# Impact of physical and cognitive exertion on cognitive control

**DOI:** 10.1101/453464

**Authors:** Karen Davranche, Gavin D Tempest, Thibault Gajdos, Rémi Radel

## Abstract

In a recent study, the differential effects of prolonged physiologically challenging 4 exercise upon two executive processes (cognitive control and working memory) have been 5 investigated. However, the impact of exercise on the selective inhibition task remained debatable and needed further analysis to dissociate the effects induced by exercise intensity from those induced by the time spent on task upon cognitive control outcomes. In this study we propose a thorough analysis of these data, using a generalized mixed model on a trial-by-9 trial basis and a new measure of the strength of the automatic response based on reaction time distribution, to disentangle the effect of physical fatigue from cognitive fatigue. Despite the prolonged duration of exercise, no decline in cognitive performance was found in response to physical fatigue. The only change observed over the 60-min exercise was an acceleration of the correct trials and an increase of errors for incompatible trials. This pattern, similar in both exercise conditions, supports the occurrence of cognitive fatigue induced by the repetition of the cognitive tasks over time.

## Impact of Physical and Cognitive Exertion on Cognitive Control

Cognitive control is essential to psychological functioning, allowing individuals to have flexible and goal-directed behaviors. In the last decade, many studies have been conducted to identify how cognitive control may be affected by physical exercise [e.g.,(Davranche et al., 2009, 2015; Joyce et al., 2009; Schmit et al., 2015)]. However, the findings are mixed as some studies have reported a decrease (Davranche et al., 2009; Davranche and McMorris, 2009; Labelle et al., 2013; Pontifex and Hillman, 2007) and others a preservation or an improvement in cognitive control during exercise (Davranche et al., 2015; Drollette et al., 2012; Schmit et al., 2015). The duration and intensity of exercise and the nature of the task have been put forward to explain the heterogeneity of the findings reported in the literature (Davranche et al., 2015; Dietrich and Audiffren, 2011). In this context, Tempest et al. (Tempest et al., 2017) tracked the dynamics of executive performance using two different cognitive tasks during cycling-exercise over a prolonged duration (60- min). While exercising at a high (physiologically challenging) or very low intensity (control condition), participants were required to perform ten blocks consisting of a cognitive control task (Eriksen task), a working memory task (n-back) and a no-task period (i.e. with no explicit cognitive demand). Each task period lasted two-minutes. The results highlighted that, compared to the very low intensity, physiologically challenging high exercise impaired working memory performance over time. However, during the cognitive control task, the interaction between exercise intensity and time approached the level of significance (p = 0.07) on mean accuracy, which made the interpretation of the impact of high exercise on cognitive control complicated. This finding indeed suggests that high intensity exercise may have impaired cognitive control efficiency over time, however a possible influence of boredom induced by the time spent on tasks could not be excluded. As suggested by the authors, this point would need to be clarified through a more extensive analysis to dissociate the potential effect of physical fatigue from the effect of cognitive fatigue upon cognitive control outcomes.

The aim of the present study was to offer this more detailed analysis of Tempest et al.’s data. Our objective was fueled by two reasons. Firstly, we wanted to reanalyse the data with a trial-by-trial analysis using a mixed model approach in order to benefit from a realistic representation of the data and to have as much information as possible (Speelman and McGann, 2013). Mixed model also provides a reduction in Type-1 (Boisgontier and Cheval, 2016) and Type-2 errors (Ma et al., 2012) in comparison to standard repeated measures analysis of variance. While Tempest and collaborators followed this recommendation and used a linear mixed model (LMM) to analyse their Eriksen task data, the skewed nature of the data exposed them to another problem. The use of LMM is conditional to the normality of the data, but dependent variables like RT are typically skewed. Tempest and collaborators therefore used the Box–Cox method (Box and Cox, 1964) to identify a transformation to normalize the data. However, this procedure can affect the magnitude (and in turn significance) of the estimated effects and complicates the interpretation of the effects. In line with prior recommendations (Dixon, 2008; Lo and Andrews, 2015; Neal and Simons, 2007), we instead fit the statistical model to the real distribution of the data by using generalized linear mixed model (GLMM), an extension of a LMM to a collection of other than normal distributions.

Secondly, and most importantly, we wanted to include distribution analysis to provide more information about the cognitive processes behind the effects found in the general analysis of RT and accuracy. Although mean RT and error rate do provide valuable information relative to cognitive processes, considering distribution analysis provides much more specific information compared to global measures classically used in most studies (van den Wildenberg et al., 2010). According to dual-route models of information processing, a conflict task such as the Eriksen task induces a conflict between an automatic and rapid response (triggered by irrelevant cues) and a slower, deliberately controlled response based on pertinent information (triggered by relevant cues). In other words, by differentiating the rapid from the slow responses, we aim to dissociate the strength of the triggering of an automatic response from the ability to inhibit proponent responses.

In the context of the study by Tempest et al. (2017), we wanted to examine how physical fatigue (continuous exertion of physical effort in the high exercise intensity condition) affects the strength of the automatic response and response inhibition. In doing so, we wanted to account for the potential effects of cognitive fatigue (induced by the repeated execution of the cognitive tasks in both the low and high intensity conditions) that could also influence automatic responses and response inhibitory processes (Faber et al., 2012; Lorist et al., 2005).

## Method

### Participants

Fourteen participants (men = 9) were recruited from a student population. Participants reported to be involved on average of 5.1 (±3.6) hours of physical activity per week. Due to the physiologically challenging intensity and duration of exercise, active participants were recruited so that they would be able to maintain the required intensity for the full 60 min (for details concerning participants’ characteristics see Tempest et al., 2017). The study was reviewed and approved by the Ethics Committee of the Université of Nice Sophia-Antipolis (*Commission scientifique de l’UFR STAPS de l’Université de Nice Sophia-Antipolis*) prior to participation and received course credits upon completion of the study.

### Procedure

This study employed a cross over design and required the participants to visit the laboratory on three occasions (one training and two experimental sessions) at least 48 h apart and around the same time of day. In the training session, participants provided their informed consent and initial assessments (age, height and body mass) were recorded. Participants were seated on an upright cycle ergometer (Wattbike, Wattbike Ltd, Nottingham, UK), previously validated for power output ranging from 50 to 300 watts (W) at a cadence of 70–90 repetitions per minute (rpm) (Hopker et al., 2010), and fitted with a facemask to measure metabolic data (FitmatePro, COSMED, Miami, USA) and a heart rate monitor. Participants completed an incremental cycling exercise test to exhaustion with an increase of 15 to 25 W per minute and a cycling cadence ranging from 70 to 90 rpm in line with the American College of Sports Medicine (Medicine, 2013). The end of the test was determined by volitional cessation of exercise or failure to maintain pedal cadence above 60 rpm despite strong verbal encouragement. Maximal oxygen uptake (VO_2_max) was determined by the highest 30 s average of oxygen uptake (VO_2_, measured in ml kg^−1^ min^−1^). The ventilatory threshold was identified by agreement of the point at which a disproportionate increase in VO_2_ occurred between two plots, (i) VO_2_ and (ii) ventilatory equivalent, over time (Gaskill et al., 2001). Participants completed a minimum of four sets (100 trials) each of the flanker and the 2-back tasks for familiarization. In order to minimize potential learning effects, additional sets of the tasks were completed until there was a <5% increase in performance from the previous set.

The order of the experimental sessions (high or very low intensity) and presentation of the tasks (flanker, 2-back or 2-back, flanker) was counterbalanced and participants were alternately assigned an order upon enrolment in the study. In the experimental sessions, the participants were seated comfortably on a cycle ergometer (SRM Trainer, SRM, Julich, Ger- many) which included supports for the forearms and two thumb response buttons on the right and left handle grips. A computer was placed at eye level in front of the participant at a distance of 80 cm. The participants then began exercise at either a workload corresponding to 10% above the ventilatory threshold (165 ± 44 W; high intensity condition) or at a very low intensity (<30 W; low intensity condition), for 60 min. The participants performed ten blocks (each lasting six minutes) consisting of the no-task (two minutes) followed by a modified version of the Eriksen flanker task and 2-back task trials (each lasting two minutes, presented in a counterbalanced order). Only the new data analysis strategy of the Eriksen task data is reported in the following paragraphs.

### Cognitive Control Task

The task consisted of a modified version of the Eriksen task (Schmit et al., 2015). In this task, each trial began with the presentation of a cross in the center of the screen as a fixation point. After 340 ms, arrow stimuli were presented in either a vertical line of three arrows or a square of 49 arrows (arranged in a 7 × 7 matrix). Each arrow on the screen was displayed as a 26 × 20 pixel symbol in black ink. Participants had to respond according to the direction shown by the central arrow. Two types of trials occurred: congruent trials (50%, all arrows pointing in the same direction) and incongruent trials (50%, center arrow pointing in the opposite direction to the other arrows). Answers were provided by pressing buttons fixed to the handlebar of the ergometer. Depending on the direction of the target arrow, the left or right button was pressed with the corresponding thumb. The delivery of the response ended the trial. If participants failed to respond within 1500 ms the trial terminated, and the next trial began immediately after. In this version, each group of arrows could randomly be displayed either at the top or at the bottom of the screen. This modification ensured a higher processing of the flankers as participants could not anticipate the location of the central arrow. The number of trials during the two minutes task was variable as it depended on the participant’s average RT.

### Data Analysis

**Reaction time.** The first trial of each block was disregarded as they quite often led to abnormally long RT due to the adjustments made by the participants to get ready to respond to the task. Decision errors and omissions were also excluded (Salthouse and Hedden, 2002). A total of 32,223 trials (87.0%) were left for further analysis. Considering that distributions of RT are positively skewed (Harald Baayen and Milin, 2010; Ratcliff, 1979), a GLMM modelled for gamma distribution (with an identity link) was used (Lo and Andrews, 2015). A random intercept effect structured by subjects was included to control for the non-independence of the data. The trial number within a block, the type of trials (congruent or incongruent trials), condition (low or high intensity), time (block 1 to 10), and all interaction between the type of trials, condition, and time were entered as fixed factors.

**Delta plot.** RT distribution analyses were obtained, separately for incongruent (IN) and congruent (CO) correct trials, using individual RT-distributions ‘‘Vincentized’’ (Ratcliff, 1979) into five equal-size speed bins (quintile). Delta plot curves were constructed by plotting the interference effect (i.e., the difference between mean RT of IN trials and mean RT of CO trials) as a function of the response speed. The data presented are the mean values of each quintile averaged across participants. Then, we examined the evolution of the magnitude of the interference as a function of the response speed. Statistical analysis was performed on the mean value of each quintile. As these values are usually normally distributed (e.g., Schmit et al., 2015) LMMs were used. A random intercept effect structured by subjects was included to control for the non-independence of the data. The order of the experimental sessions (to control for order effects within the data), condition (low or high intensity), time (block 1 to 10) and quintile factor (from 1 to 5) were entered as fixed factors. Another statistical analysis was also performed on the slope of the last segment which is routinely used to measure the efficiency of response inhibition [see, van den Wildenberg et al., 2010].

**Accuracy.** After excluding omissions and the first trial of each block, a total of 36611 (98.9%) trials were left in the analysis. Since the dependent variable is a dichotomous variable (0=errors; 1=correct responses), it can be easily modelled using a GLMM for a logistic distribution (Dixon, 2008). A random intercept effect structured by subjects was included to control for the non-independence of the data. The trial number within a block, the type of trials (CO or IN trials), condition (low or high intensity), time (block 1 to 10), and all interactions possible between the type of trials, condition, and time were entered as fixed factors.

**Error Location Function and Error Location Index.** We assessed the effect of physical and cognitive fatigue on the strength of the automatic response using a new measure of the strength of the automatic response recently proposed by Servant, Gajdos, and Davranche (2018): the Error Location Function (ELF), which represents the proportion of errors located below each quantile of the overall RT distribution, in the first three and the last three blocks performed in both exercise conditions. Specifically, we quantified the strength of the automatic response by the Error Location Index (ELI), which is equal to the area under the ELF curve. The ELI can be interpreted as the expectation that a uniformly drawn incorrect response is faster than a uniformly drawn (correct or incorrect) trial. Thus, if ELI = 1, all errors are concentrated among the fastest trials, which corresponds to a very strong response capture. On the other hand, if ELI = 0, all errors are concentrated among the slowest trials, which is the converse of response capture. In general, a higher ELI indicates a strongest response capture. A LMM was performed on ELI values to assess the effect of time on task and the effect of exercise intensity on the automatic response. The condition (low or high intensity) and time (block 1 to 10) were entered as fixed factors.

For all statistical models, it is important to note that while block was entered as a linear covariate in Tempest et al.’s study, it was entered as a categorical factor in the present models. This choice was made for two reasons. First, we believe that the additional free parameters related to the inclusion of this variable as a categorical factor is not a problem considering the greater number of trials. In addition, entering a variable as a covariate assumes that the effect is linear and constant over time. However, our hypotheses related to fatigue suggests that the effects should be localized in the last blocks. All statistics were performed using SPSS (version 23, IBM statistics, Armonk, NY, USA). Significant main and interaction effects were reported (*p*<.05) and followed-up using Sidak-adjusted multiple comparisons tests. To report descriptive statistics, we provided the means ± standard errors estimated by the models to represent the effect of interest after controlling for other factors and covariates.

## Results

### Reaction Time

The GLMM revealed a significant compatibility effect [F(32461, 1)=2152.312, p<.001], indicating faster RT for CO (465.0±8.4ms) than IN trials (530.3±9.6ms). A condition effect was found [*F*(32461, 1)= 125.69, *p*<.001], indicating faster RT in the high (488.7±8.9ms) than low intensity exercise (504.6±9.2ms) condition. A time effect was found [*F*(32461, 9)=8.13, *p*<.001], indicating slower RT in the first two blocks than in the following blocks (block 1 was significantly different from block 3, 4, 5, 6, 7, 8, 9, and 10 and block 2 was significantly different from block 6, 7, 9, and 10). In addition a significant interaction between block and condition was found [*F*(32461, 1)=2152.31, *p*<.001], indicating that the two exercise conditions were only significantly different at block 2, 3, 4, 6, 7, 8, 9 and 10 (Figure 1). Neither the interaction between block and compatibility [*F*(32461, 9)= 0.48, *p* = .89], nor the interaction between condition and compatibility [*F*(32461, 1)= 1.09, *p =*.30] or between these three factors [*F*(32461, 9)= 0.47, *p =* .89] were significant.

**Figure 1.**
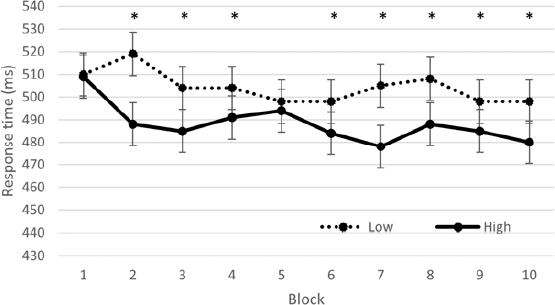
Reaction time (ms) on the Eriksen task, as a function of time on task (block number) and condition (low vs. high intensity). * indicates a significant difference (p<.05) between the low and high condition.

### Accuracy

The GLMM revealed a significant effect of type of trials [*F*(36079, 1)=15.05, *p*<.001] indicating a linear increase of the number of errors within a block (coefficient=.002). A significant effect of block was also found [*F*(36079, 9)=3.17, *p*=.001] indicating that the number of errors were significantly larger in block 9 than in the block 2, 3, and 4. A compatibility effect was found [*F*(36079, 1)=1615.09, *p*<.001] with more errors for IN (15.5±2.5%) than for CO (2.7±0.5% of errors) trials. A condition effect was found [*F*(36079, 1)=28.49, *p*<.001] with more errors in the high (7.5±1.3%) than low (5.9±1.1% of errors) exercise conditions. An interaction between compatibility and block was also found [*F*(36079, 1)=3.23, *p*<.001], indicating that while accuracy did not evolve over time for CO trials, it led to an overall increase of errors over time for IN trials. For the IN trials, block 1 was significantly different from block 5, 6, 9, and 10; block 2 was significantly different from block 5, 9, and 10; and block 3 and 4 were significantly different from block 9 and 10 (see Figure 2). Neither the interaction between condition and block [F(36079, 9)=0.73, *p*=.68], nor the interaction between condition and compatibility [F(36079, 1)=0.38, *p*=.54] or between these three factors [F(36079, 9)=0.40, *p*=.94] were significant.

**Figure 2.**
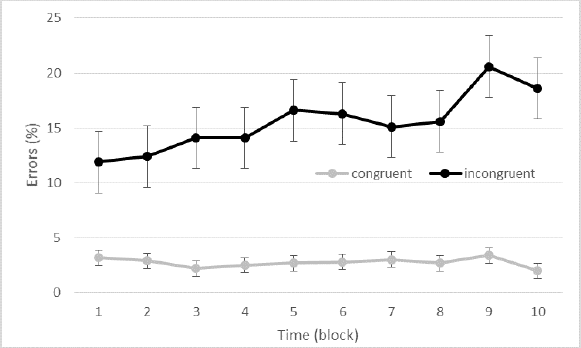
Accuracy (percent of errors) as a function of time and congruency (congruent vs. incongruent trials).

### Delta Plots

The first LMM including the mean values of each quintile revealed a condition effect [*F*(1, 1287)=6.25, *p*<.02], indicating smaller interference during exercise in the high (64.597 ± 4.085ms) than in the low intensity exercise (69.795 ± 4.085ms) condition. Beside the logical quintile effect [*F*(1287, 4)= 12.79, *p* < .001], neither the interaction between block and quintile [*F*(1287, 36)= 0.471, *p* = 1], nor the interaction between condition and quintile [*F*(1287, 4)= 0.201, *p =*.94] or between these three factors [*F*(1287, 36)= 0.28, *p =* 1] were significant.

The second LMM performed on the slope of the last segment revealed no significant effect. Neither the effect of block [*F*(247, 9)=0.66, *p*=.74], nor the effect of condition [*F*(247, 1)=0.53, *p*=.47] or the interaction between these two factors [*F*(247, 9)=0.54, *p*=.84] were significant.

### Error Location Index

The LMM analysis including the ELI of the first three blocks and the last three blocks performed in both exercise conditions revealed a block effect [*F*(39, 1)=4.43, *p*=.04] indicating a decrease of the ELI index in the terminal blocks (.749 ± .018) compared to the initial blocks (.776 ± .018, see Figure 3). Neither the effect of condition [*F*(39, 1)=0.94, *p*=.34], nor the interaction between block and condition [F(39, 1)=0.004, *p*=.95] were significant. The decrease of ELI index indicated a lower concentration of errors for IN trials among the fastest trials in the terminal blocks, i.e., that the strength of the automatic response was lower over time.

**Figure 3.**
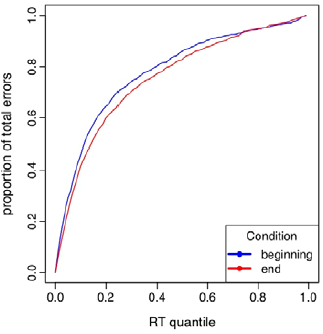
Error location function (ELF) curves aggregated across subjects in the first three blocks (beginning; blue) and the last three blocks (end; red). Error location index represents the area under ELF curves.

## Discussion

The objective of the present research was to offer a thorough analysis of Tempest et al.’s data which tracked changes of higher-order cognitive performance during 60-minute of prolonged cycling-exercise. The aim of the present study was to assess the impact of prolonged high exercise on cognitive control (i.e., Eriksen task) and to dissociate the effect of physical fatigue induced by exercise from cognitive fatigue induced by the repetition of the cognitive tasks over time.

By investigating changes in cognitive functioning through trial-by-trial and distributional analyzes, the present study confirmed a speed-up of both congruent and incongruent trials in the high condition compared to the low exercise intensity condition. This benefit is not in discrepancy with the current literature on exercise and cognition and could be explained by an increase in the brain activation level globally improving the information processing at both the sensory and motor levels (Davranche et al. 2005; 2006). Even if participants reported working very hard in the terminal blocks of the high intensity condition, RT performance were enhanced and we did not observe deterioration of cognitive control induced by physical fatigue over the 60 min-exercise. Instead, the analysis performed on the magnitude of the interference highlighted a smaller interference effect during exercise at a high intensity than at a low intensity. In other words, physiologically challenging exercise also enhances performance in a conflict task requiring the participant to inhibit the automatic response triggered by task-irrelevant aspect of the stimulus. This finding suggests that exercise could enhance at least part of online control mechanisms. It is noteworthy that, as already reported by Schmit et al. (2015), the facilitating effect usually reported during moderate exercise appears to also occur during intense and prolonged exercise. Response inhibition did not seem to be affected by the time spent on the task. Whatever the intensity of exercise, the magnitude of the interference remained constant over time. This finding does not appear surprising since proficient cognitive control has already been reported during intense physical exercise (Davranche et al., 2015; McMorris et al., 2009), physical exercise maintained until exhaustion (Schmit et al., 2015) and after complete sleep deprivation (Temesi et al., 2013).

In contrast, RT for both type of trials (CO and IN) were faster and results showed an overall increase of errors for IN trials over time. The repetition of the cognitive tasks over 60- min with ten 2-min blocks of an Eriksen task and ten 2-min blocks of a working memory task altered performance on the conflict task for both exercise intensity conditions. This observation can be assimilated to the large literature showing that executive performance tends to decrease over time (e.g., Faber et al., 2012; Lorist et al., 2005; Persson, Welsh, Jonides, & Reuter-Lorenz, 2007). While this effect has been demonstrated in the context prolonged continuous performance (Faber et al., 2012), it has also often been shown in the context of repeated performance (Dang, 2017). While these effects have been typically ascribed to cognitive fatigue (Muraven and Baumeister, 2000), it is actually very difficult to identify the true origins of this effect, as both (loss of) motivation and fatigue can be advanced to explain the decline of cognitive performance in time (Boksem et al., 2006; Boucher and Kofos, 2012; Inzlicht and Schmeichel, 2012).

This decreased accuracy, which appeared with the lengthening of the time spent on task in very low intensity as in high intensity, probably results from a complex state involving multiple changes such as modifications of motivation, cognition, and mood. With the occurrence of this fatigue-related decrement in performance, it would make sense to speculate that participants may, voluntarily or not, reduced their task engagement and opt for an easier strategy leading them to progressively act based on impulsive automatic responses rather than based on supervised and controlled responses. If this had been the case, we should have observed an increase of the ELI index due to an increase of the relative proportion of fast errors, which typically corresponds to an increase of the strength of automatic response. The current results highlighted the opposite effect, showing a decrease of the ELI index which indicates a lower concentration of errors for IN trials among the fastest trials in the terminal blocks.

To sum up, the present reanalysis of Tempest et al.’s data allowed dissociating the effect of exercise intensity, the effect of physical fatigue and the effect of cognitive fatigue. The main finding is that, compared to the very low intensity, physiologically challenging high exercise did not alter cognitive control and rather enhanced performance in a conflict task involving response inhibition. In the context of an inconsistent literature concerning the impact of exercise on executive functions, the present findings add a further argument in favor of the robustness of the cognitive control. Despite the prolonged duration of the high exercise for 60min, no decline in cognitive performance induced by physical fatigue was found, even in the terminal blocks. The only change observed in cognitive performance over the 60-min was a shortening of overall RT and an increase of errors for IN trials. The fact that these findings were similar for both exercise intensity conditions, supports the idea of the occurrence of cognitive fatigue with the repetition of the cognitive tasks over time. The present study provides important new insights regarding the specific changes in cognitive processes during intense prolonged exercise and opens new perspectives regarding field assessment of cognitive mechanisms underlying cognitive fatigue. In this regard, considering the whole distribution of every trial using a GLMM is a useful approach which provides a relevant statistical model and a detailed analysis to assess changes in cognitive functioning.

## Acknowledgments

This work is supported by a public grant overseen by the French National Research Agency (ANR) [reference: ANR-13-JSH2-0007]. The results of this study are presented clearly, honestly, and without fabrication, falsification, or inappropriate data manipulation.

Contributions: RR, GDT and KD designed the study; RR and GDT carried out the experiment; RR, KD and TG analysed the data and wrote the manuscript; RR supervised the project.

Conflict of Interest: The authors have no conflict of interest to declare.

